# Far-red light enhances soluble sugar levels and *Botrytis cinerea* disease development in tomato leaves in a jasmonate-dependent manner

**DOI:** 10.1101/2020.05.25.114439

**Authors:** Sarah Courbier, Sanne Grevink, Emma Sluijs, Pierre-Olivier Bonhomme, Kaisa Kajala, Saskia C.M. Van Wees, Ronald Pierik

**Author notes:** **Author correspondence:** Ronald Pierik.

## Abstract

Plants lacking phytochrome photoreceptors display elevated soluble sugar levels in leaves. Although pathogens principally feed on sugars supplied by the plant, the link between increased plant sugar levels upon phytochrome inactivation and disease development has not been considered.

Tomato plants were exposed to control white LED (WL) or a combination of white and far-red LED (WL+FR) light, to inactivate phytochrome signaling and modulate soluble sugar levels. We also experimentally manipulated internal sugar levels by either supplementing glucose or inhibiting photosynthesis in tomato leaflets prior to performing soluble sugar quantifications or bioassays with pathogens.

Tomato plants exposed to WL+FR or lacking phytochrome B (*phyB1phyB2* double mutants) show enhanced levels of soluble sugars, especially glucose and fructose, in their leaves. The jasmonic acid biosynthesis mutant *def1* also has elevated soluble sugar levels, which could be rescued by exogenous methyl-jasmonate application. This indicates an interplay between JA signaling and primary metabolism.

The increase in soluble sugar levels in tomato leaves upon phytochrome inactivation is regulated in a JA-dependent manner. Our data stress the importance of primary metabolism in the FR-induced susceptibility in tomato that could contribute to promote plant resistance when grown at high density.

## Introduction

As photoautotrophic organisms, plants harness light energy into sugars through photosynthesis. Red (R) and blue (B) light are absorbed by plant tissue while far-red (FR) light is reflected by or transmitted towards neighboring vegetation. At high planting density, plants experience a drop in the R:FR ratio due to the depletion of R and the omnipresence of FR radiation in the canopy. Changes in the R:FR ratio in the environment are sensed by specialized photoreceptors called phytochromes, where phyB plays a dominant role and can be found in two photoconvertible forms, the active (Pfr) and inactive (Pr) form (Franklin, 2008). The perception of low R:FR conditions subsequently leads to photoconversion of phyB into the inactive Pr form that is restricted to the cytoplasm. This phenomenon prevents phyB interaction with the bHLH-type Phytochrome Interacting Factors (PIF) transcription factors and their consequent inactivation and degradation by the 26S proteasome (Li *et al.*, 2016). The resulting accumulation of PIFs initiates downstream growth responses, largely via increased auxin synthesis and response in Arabidopsis known as shade avoidance (Hornitschek *et al.*, 2012; Li *et al.*, 2012; Pantazopoulou *et al.*, 2017). The perception of low R:FR or the genetic inactivation of phyB in plants is known to affect plant immunity towards distinct pathogens (Ballaré, 2014; Ballaré & Pierik, 2017, Pieterse *et al*., 2014). This so-called FR-induced susceptibility is an example of the well-known growth-defense trade-off, and has been studied in a number of plants species and pathosystems. Phytochrome inactivation upon low R:FR promotes lesion development induced by the necrotrophic fungus *Botrytis cinerea* and enhances *Mamestra brassicae* and thrips feeding performance in Arabidopsis and tomato (Cortés *et al.*, 2016; Izaguirre *et al.*, 2006; Ji *et al.*, 2019). Compromised plant defense towards *B. cinerea* in low R:FR conditions goes via a decrease in secondary metabolite production and defense gene activation in Arabidopsis (Cargnel *et al.*, 2014; De Wit *et al.*, 2013). Interestingly, these are not passive trade-offs, but involve molecular control. During FR-enrichment, the gibberellin (GA) pathway is strongly activated to promote shoot elongation via degradation of growth-inhibiting DELLA proteins. However, DELLA proteins not only inhibit growth, but also promote jasmonic acid (JA) signaling by physically interacting with JAZ proteins, negative defense regulators (Cerrudo *et al.*, 2012; Leone *et al.*, 2014, Hou *et al.*, 2010). DELLA removal in low R:FR therefore releases JAZ proteins that in turn sequester MYC transcription factors, thereby preventing the activation of JA-associated defense gene expression (Leone *et al.*, 2014; Pieterse *et al.*, 2014). In addition, low R:FR conditions were also shown to reduce the pool of bioactive JAs available in Arabidopsis plant tissue upon exogenous MeJA treatment and was associated with increased susceptibility to *B. cinerea* (Fernández-Milmanda *et al.*, 2020). Although the DELLA-JAZ interaction or the decrease in JA response explain part of the FR-induced susceptibility, there likely are several other layers of regulation involved in balancing shade avoidance and defense against pathogens and pests.

In addition to modulating plant defense against pathogens, phytochrome signaling has also been described to influence carbohydrate levels in plants. In Arabidopsis, plants lacking phytochromes showed an increase in soluble sugars and starch during daytime compared to wild-type plants indicating the involvement of phytochrome in the control of primary metabolism (Yang *et al.*, 2016). In addition, low R:FR treatment promotes carbohydrate levels various species, including *Brassica rapa* (De Wit *et al.*, 2018), *Allium cepa* (Lercari, 1982), *Cucumis sativus* (Xiong *et al.*, 2011) and *Nicotiana tabacum* (Kasperbauer *et al.*, 1970). Carbohydrates generated by plants are also the primary nutrient source for plant pathogens. It is known that plants strictly regulate sugar fluxes to restrict pathogens from accessing those resources. However, plant pathogens have developed the capacity to hijack the sugar transport machinery of their host to redirect carbohydrates towards the infection site (Chen *et al.*, 2010; Doidy *et al.*, 2012; Lapin & Van den Ackerveken, 2013). For instance, pathogenic bacteria and fungi were shown to promote the expression of different sets of SWEET sugar transporters in Arabidopsis upon infection showing a pathogen-specific sugar transport manipulation (Chen *et al.*, 2010). In grapevine, *B. cinerea* has been shown to upregulate *VvSWEET4* and further analysis on Arabidopsis showed that *sweet4* mutants are less vulnerable to the fungus (Chong *et al.*, 2014). Although these studies mostly focus on the impact of a pathogen infection on the sugar allocation of the host, the impact of enhanced soluble sugar levels (e.g. via phytochrome inactivation) on disease severity has not received much attention. High sugar concentrations in plant tissue, although beneficial for the pathogen, are often associated with stronger plant defense potential. Studies have shown the involvement of sugars in the induction of secondary metabolites such as phenylpropanoids and/or defense gene induction such as MAPK or PR genes (reviewed in Moghaddam & Van Den Ende, 2012). In addition, a negative correlation between the defense hormone JA and soluble sugar levels was observed in *Nicotiana attenuata* (Machado *et al.*, 2015). JA biosynthesis mutant plants displayed elevated monosaccharide levels in leaf tissue as well as increased *Manduca sexta* larval mass upon infestation compared to wild-type plants (Machado *et al.*, 2015). Also, a recent study reported that the disease severity induced by *B. cinerea* is negatively correlated to the relative fructose content in tomato stem (Lecompte *et al.*, 2017). Nevertheless, JA and sugar signaling pathways are not always negatively correlated. Studies in Arabidopsis have shown that glucose and JA can act synergistically to increase glucosinolate (GS) levels which have been shown to promote plant resistance towards *B. cinerea* in Arabidopsis (Buxdorf *et al*., 2013; Guo *et al.*, 2013; Kliebenstein *et al.*, 2005). Overall, even though it becomes clear that plant soluble sugar status can influence plant immunity, it is still unclear whether elevated soluble sugar levels would promote or inhibit disease development.

Here, we propose that phyB inactivation could enhance disease severity via soluble sugar accumulation in plant tissue. We show that tomato plants exposed to WL+FR or lacking phyB have increased levels of glucose and fructose in leaves compared to WL-treated plants. By experimentally modulating soluble sugar levels in tomato leaves, we associate high soluble sugar levels to enhanced symptom development induced by *B. cinerea*. Furthermore, a tomato mutant deficient in JA biosynthesis exhibited high soluble sugar levels in leaf tissue accompanied with elevated disease severity. Consistent with these findings, soluble sugar levels as well as plant susceptibility was diminished by exogenous MeJA both in the JA-biosynthesis mutant and wild-type plants showing the occurrence of JA-mediated soluble sugar modulation in tomato leaves. As JA responsiveness is decreased in tomato phytochrome mutants (Cortés *et al.*, 2016), we propose that the elevated soluble sugar levels found in WL+FR-treated plants can be mediated by reduced JA signaling in low R:FR light. These results indicate that the FR-induced susceptibility in tomato leaves is partly controlled by phyB signaling as it regulates JA-mediated soluble sugar modulation in turn affecting disease severity induced by pathogens.

## Material and methods

### Plant growth and light treatments

Seeds from tomato (*Solanum lycopersicum*) cv. Moneymaker and *phyB1phyB2* double mutant (cv. Moneymaker), curated within the LED it Be 50% consortium, as well as tomato cv. Castlemart and JA biosynthesis mutant *defenseless 1* (*def1*) (cv. Castlemart), kindly provided by Dr. Maria J. Pozo (Dept Soil Microbiology and Symbiotic Systems, Granada, Spain), were sown in wet vermiculite. After 10 days, tomato seedlings were transferred into 9 × 9 cm pots with regular potting soil (Primasta® soil, the Netherlands). Plants were grown for four weeks after sowing in climate chambers (MD1400; Snijders, the Netherlands) in long-day photoperiod (8-h dark / 16-h light) at 150 μmol m^−2^ s^−1^ photosynthetically active radiation (PAR) using white light (WL, R:FR ratio = 5.5) generated by Philips GreenPower LED research modules (Signify B.V.). The calculated phytochrome photostationary state (PSS) value was 0.8 equivalent to the proportion of active phytochrome (Pfr) within the total of active and inactive (Pr) phytochrome present (Sager *et al.*, 1988). For additional FR treatments, three-week-old plants were exposed for five days, starting at Zeitgeber time (ZT) = 3 on the first day, to WL supplemented with uniform far-red (FR) Philips GreenPower LED research modules referred to as WL+FR (R:FR = 0.14, PSS value = 0.5). Local FR radiation was performed by using custom FR LEDs (724-732 nm) attached onto a flexible arm (Fig. **S1**). The top leaflet of either the 3^rd^ or 4^th^ oldest leaf of four-week-old WL-exposed plants were illuminated with FR. WL-exposed plants which did not receive local FR illumination were taken as control.

### Pathogen growth conditions and bioassays

#### Botrytis cinerea

*B. cinerea* strain *Bc* 05.10 (Van Kan *et al.*, 1997) and a pepper isolate (Denby *et al.*, 2004; Grant *et al.*, 2003) were maintained on half strength Potato dextrose agar medium (PDA ½, BD Difco^TM^) and grown for approximately 10 days at room temperature under natural daylight conditions. The spore suspensions were prepared according to Van Wees *et al.* (2013) and diluted to a final concentration of 1.5 × 10^5^ spores ml^−1^ in half strength potato dextrose broth (PDB ½, BD Difco™) prior to inoculation. Bioassays were performed on detached tomato leaflets previously pretreated with WL or WL+FR for five days. The leaflets were placed in square Petri dishes onto Whatman® filter soaked with 6 ml of tap water to avoid dehydration. The adaxial side of the leaflets was drop-inoculated 3-6 times with 5 μl of spore suspension. Plates were sealed with PARAFILM® M and incubated for 3 days in their respective light pretreatment conditions (WL or WL+FR). Pictures were taken 3 days post inoculation (dpi) and lesion areas were measured using the imageJ software.

*In vitro* mycelium diameter measurements were performed by depositing a 5 μl droplet of a 1.5 × 10^5^ spores ml^−1^ solution onto 1.5 % granulated agar (BD Difco™) plates supplemented with homogenates of WL- or WL+FR-treated leaves (1 gFW ml^−1^), or onto PDA ½ supplemented with increasing concentrations of D-glucose (Duchefa Biochemie) (10, 20 and 50 g l^−1^). Inoculated plates were incubated for three days in WL at room temperature. The newly grown mycelium diameter was measured by using a digital caliper.

#### Phytophthora infestans

*P. infestans* strain Pi88069 (Kamoun *et al.*, 1997) was maintained on Rye sucrose agar medium (RSA, Caten & Jinks, 1968) for 10 days in darkness at room temperature. Cultured plates were flooded with 10 ml of sterile MilliQ water and incubated at 4°C for 3 hours. Zoospores were collected, counted and adjusted to a concentration of 5 × 10^4^ zoospores ml^−1^ prior to the bioassays. The abaxial side of detached tomato leaflets was drop-inoculated several times with 10 μl of zoospores suspension and leaflets were incubated for 3 days in their respective light pretreatment conditions (WL or WL+FR). The lesion areas were measured after 3 dpi with the ImageJ software. Statistical analysis was performed with student t-test, p-value < 0.05.

*In vitro* mycelium diameter measurements were performed by depositing a plug of mycelium (ø 0.6 cm) in the center of ½ strength V8 agar plates (1.5 % granulated agar, BD Difco™) supplemented with increasing concentrations of D-glucose (Duchefa Biochemie) (0, 10, 20 and 50 g l^−1^). Inoculated plates were incubated for 10 days in darkness. The newly grown mycelium diameter was measured by using a digital caliper.

#### Pseudomonas syringae

*P. syringae* pathovar tomato strain DC3000 (Whalen *et al.*, 1991) was grown in liquid King’s medium B (KB) (King *et al.*, 1954) and incubated overnight in an orbital shaker at 220 rpm at 28°C. The next day, the bacterial culture was centrifuged at 7800 rpm (7170 G) for 5 min. The bacterial pellet was washed 3 times with sterile MilliQ water and resuspended in 10 mM MgSO4. The inoculum was adjusted to OD_660_ = 0.025 (OD_660_ = 1 = 10^9^ cells ml^−1^) and supplemented with 0.02 % Silwet L-77 (Lehle seeds). The two first lateral leaflets of the 3^rd^ leaf of WL- and WL+FR-pretreated intact tomato plants were dipped in the bacterial inoculum for 3 seconds and whole plants were subsequently placed in their respective light treatment for 3 days at 95% RH. Bacteria were extracted by grinding two leaf discs (ø 0.6 cm) from each leaflets in 400 μl of 10 mM MgSO_4_. Dilutions (10^−2^ to 10^−5^) of the homogenates were plated and incubated at 28°C on KB medium supplemented with 25 μg ml^−1^ rifampicin until colony forming units (cfu) appeared. Bacterial growth was calculated in cfu per cm^2^.

For glucose-mediated bacterial growth measurements were performed by growing bacteria in liquid KB until OD_660_ reached 0.1. Per replicate, 4 ml of bacterial suspension was transferred into a new tube and supplemented with 1 ml of water or with glucose solutions at different concentrations (10, 20 and 50 g l^−1^). The OD_660_ was measured every hour for 3 hours.

### *In planta* modulation of soluble sugar levels

Tomato leaflets of the 3^rd^ leaf were detached under water and placed in square Petri dishes containing Whatman® filter paper (bottom 1/3 cut out) soaked with 2.5 ml of tap water to avoid dehydration. Plates were placed on a metallic rack at a 30° angle. The bottom of the tilted plates was filled with 13 ml of 0.5 M glucose solution for glucose supplementation experiments or 100 μM 3-(3,4-dichlorophenyl)-1,1-dimethylurea (DCMU, Sigma®) for sugar starvation experiments. Tap water was used as a control in mock conditions. Only the petiole of the leaflets was immersed in the solution. Plates were kept at a 30° angle and incubated in WL or FR conditions for 24 h prior to sugar quantifications or bioassays with *B. cinerea*. Prior to inoculation, the leaflets were transferred into square Petri dishes and drop-inoculated following the procedure described above.

### Plant hormone treatments

Three-week-old tomato plants were placed under either WL or WL+FR for five days and were sprayed daily with a 50 μM methyl jasmonate (MeJA, Sigma-Aldrich®) or mock solution (0.1 % EtOH) supplemented with 0.1 % Tween20 (Duchefa biochemie). Hormone treatments were performed at 10 a.m. every day (ZT = 3). For sugar quantifications, leaf discs (ø 0.6 cm) originating from the 3^rd^ oldest leaf were punched and snap frozen in liquid nitrogen prior to performing soluble sugar extraction and quantifications. For bioassays, treated leaflets were transferred in square Petri dishes and inoculated with *B. cinerea* spores as described above.

### Extraction and quantification of soluble sugars

Tomato leaf discs (15 – 20 mg of fresh tissue) were harvested and snap frozen in liquid nitrogen. Samples were ground, supplemented with 135 μl of 0.83 M perchloric acid (HClO_4_, Merck), vortexed and centrifuged for 15 min at 4 °C and 13000 rpm using a table centrifuge (19330 G). The supernatant (110 μl) containing soluble sugars was transferred into a new tube and the pellet containing starch and proteins was kept at −80 °C for starch quantification analysis. The samples were mixed with 25 μl of 1 M Bicine (Sigma-Aldrich®) and ~22 μl 4 M KOH solution to neutralize the pH to 7. All samples were centrifuged for 10 min at 4 °C and 13000 rpm and the supernatant was collected into a new tube for sugar quantifications that were performed using ©Megazyme Sucrose/D-Fructose/D-Glucose Assay Kit (K-SUFRG, ©Megazyme) with a few modifications. Per sample, 30 μl of sugar extract or of a dilution series from 2000 μM to 0 μM glucose solution (used as standards) were mixed with 185 μl of reaction mix containing 170 μl MilliQ, 10 μL of Buffer (solution 1) and NADP+/ATP (solution 2) in a transparent flat-bottomed 96-well plate. The absorbance at 340 nm (A_340nm_) was measured at the start of the experiment as well as after a 15 min incubation at 37 °C (R_0_ and R_1_). All wells were supplemented with 2 μL of HXK/G6DPH (solution 3) prior to 15 min at 37°C and A_340nm_ measurement (R_2_). The procedure was repeated by adding 2 μL of PGI (solution 4) incubating for 15 min at 37 °C and measuring A_340nm_ (R_3_). The same incubation and A_340nm_ measurements were performed after adding 2 μL of yeast invertase solution (100 mg ml^−1^, Sigma-Aldrich®) diluted in buffer (solution 1) were added to each well and incubated for 25 min at 37 °C before the final A_340nm_ measurement (R_4_). The glucose standard curve equation (*y* = *ax* + *b*) was used for soluble sugar quantifications. Formulas used to determine soluble sugar concentrations:

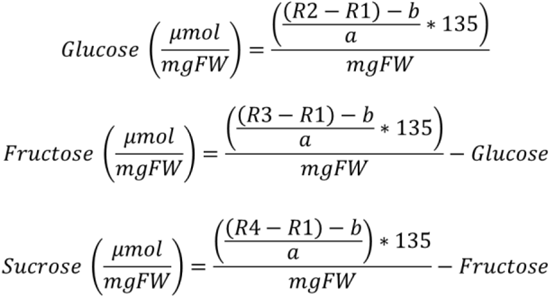

## Results and discussion

### Phytochrome inactivation enhances soluble sugar levels and disease severity of *B. Cinerea* in tomato

Previous studies have shown that Arabidopsis phytochrome mutants accumulate more soluble sugars during the day than the wild type does (Yang *et al.*, 2016). First, we studied whether, as in Arabidopsis, a *phyB* mutation would affect soluble sugar levels in tomato leaves and could in turn lead to increased disease severity towards the necrotrophic fungus *B. cinerea*. Since tomato has two *PHYB* genes, we grew the *phyB1phyB2* double mutant under white light LEDs (WL) for three weeks. We recorded stem elongation for five days prior to quantifying soluble sugar levels in leaves and inoculation with *B. cinerea* spores (Fig. **1**). Tomato *phyB1phyB2* plants exhibited a constitutive shade avoidance response through a strong stem elongation phenotype (Fig. **1a**) as well as elevated glucose and fructose levels in leaves compared to wild-type Moneymaker plants (Fig. **1b**). This was accompanied by an increased foliar susceptibility to *B. cinerea* in *phyB1phyB2* compared to wild-type Moneymaker plants (Fig. **1c**). Possibly, the increase in soluble sugars in *phyB1phyB2* mutants might benefit *B. cinerea* growth *in planta*, thereby enhancing disease development on tomato leaves.

**Figure 1:**
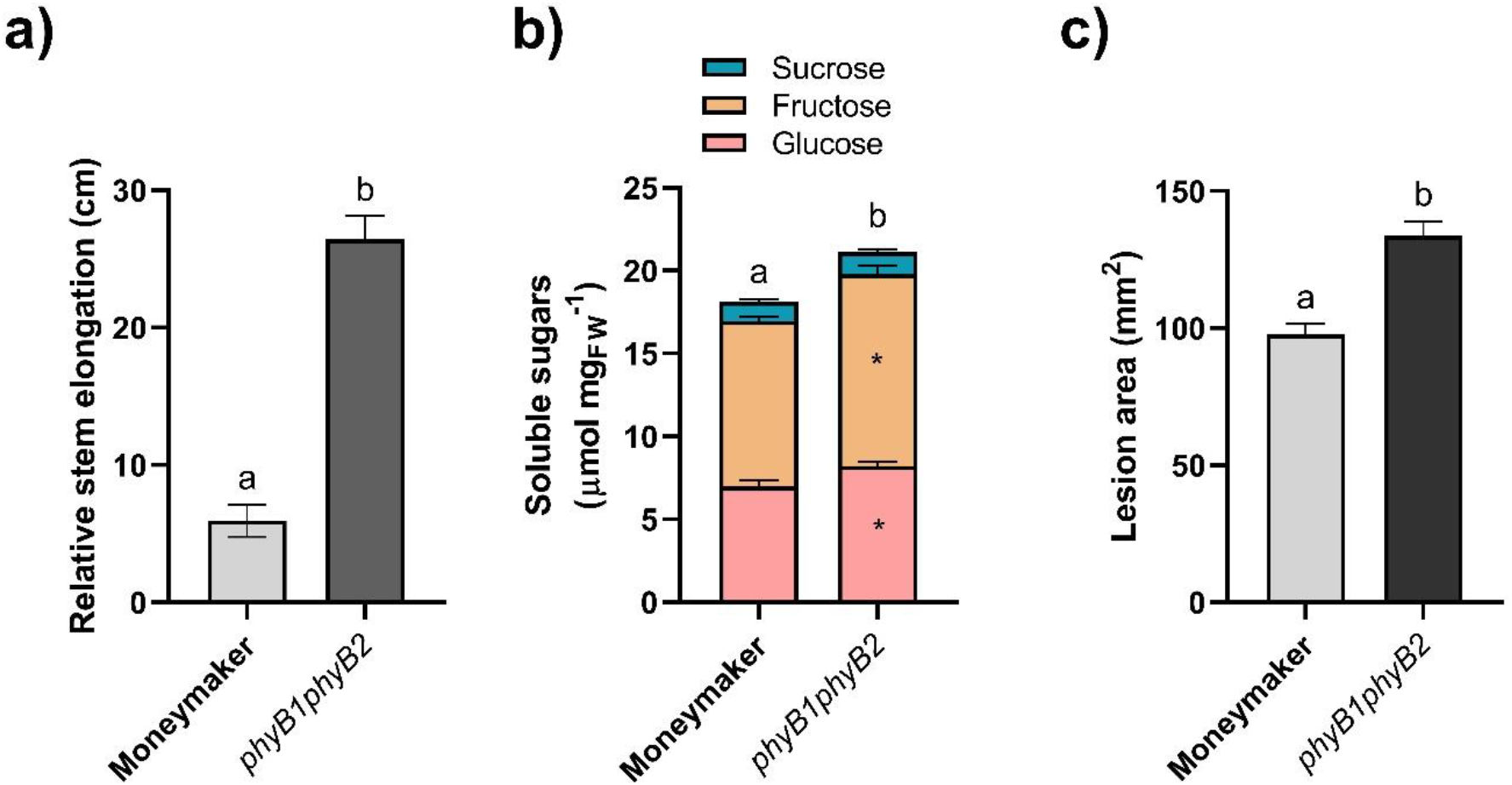
*phyB1phyB2* double mutants display increased susceptibility to *Botrytis cinerea* associated with increased soluble sugar levels in leaves. **(a)** Relative stem elongation over five days for three-week-old Moneymaker and *phyB1phyB2* mutants. Data correspond to day_5_-day_0_ measured at ZT = 3. **(b)** Soluble sugars (glucose, fructose and sucrose) quantified at midday (ZT = 8) for three-week-old WL-grown Moneymaker and *phyB1phyB2*. **(c)** Lesion area induced by *B. cinerea* spores on Moneymaker and *phyB1phyB2* detached leaflets measured at 3 dpi. Data correspond to the mean ± SEM, different letters represent significant differences based on ANOVA post-hoc test and asterisks based on Student’s t-test (p < 0.05). n = 5-10 plants per treatment.

Next, we studied whether alike the *phyB1phyB2* double mutant (Fig. **1b**), inactivating phytochrome by supplementing the WL background with FR LED light (WL+FR) would increase soluble sugar levels in leaves. Over a course of five days of light treatment, we observed that soluble sugar levels, mainly glucose and fructose, were significantly increased in WL+FR-exposed leaves compared to the WL control starting at 3 days of treatment (Fig. **2a-d**). Altogether, our data demonstrate that the increase in soluble sugar levels occurring phytochrome mutants previously shown in Arabidopsis (Yang *et al.*, 2016) is also observed in tomato (Fig. **1b** and Fig. **2a**). It seems paradoxical that WL+FR conditions would at the same time promote energy and carbon-requiring shoot elongation and soluble sugar accumulation in leaf tissue. However, since we measured sugars in tomato leaflets specifically, we cannot rule out the possibility that soluble sugars would accumulate in leaves at the expense of other plant organs. We hypothesize that WL+FR would cause a gradual sugar accumulation in leaf tissue beyond levels that would be fully consumed in the dark period leading to leftover carbohydrates in the leaves from the third day of treatment onwards (Fig. **2a**). Indeed, we could only observe an increase in susceptibility when leaves had been exposed to WL+FR for at least four days (Fig. **2e**), consistent with the proposition that FR-induced sugar accumulation in tomato leaves could promote *B. cinerea* development in plant tissue.

**Figure 2:**
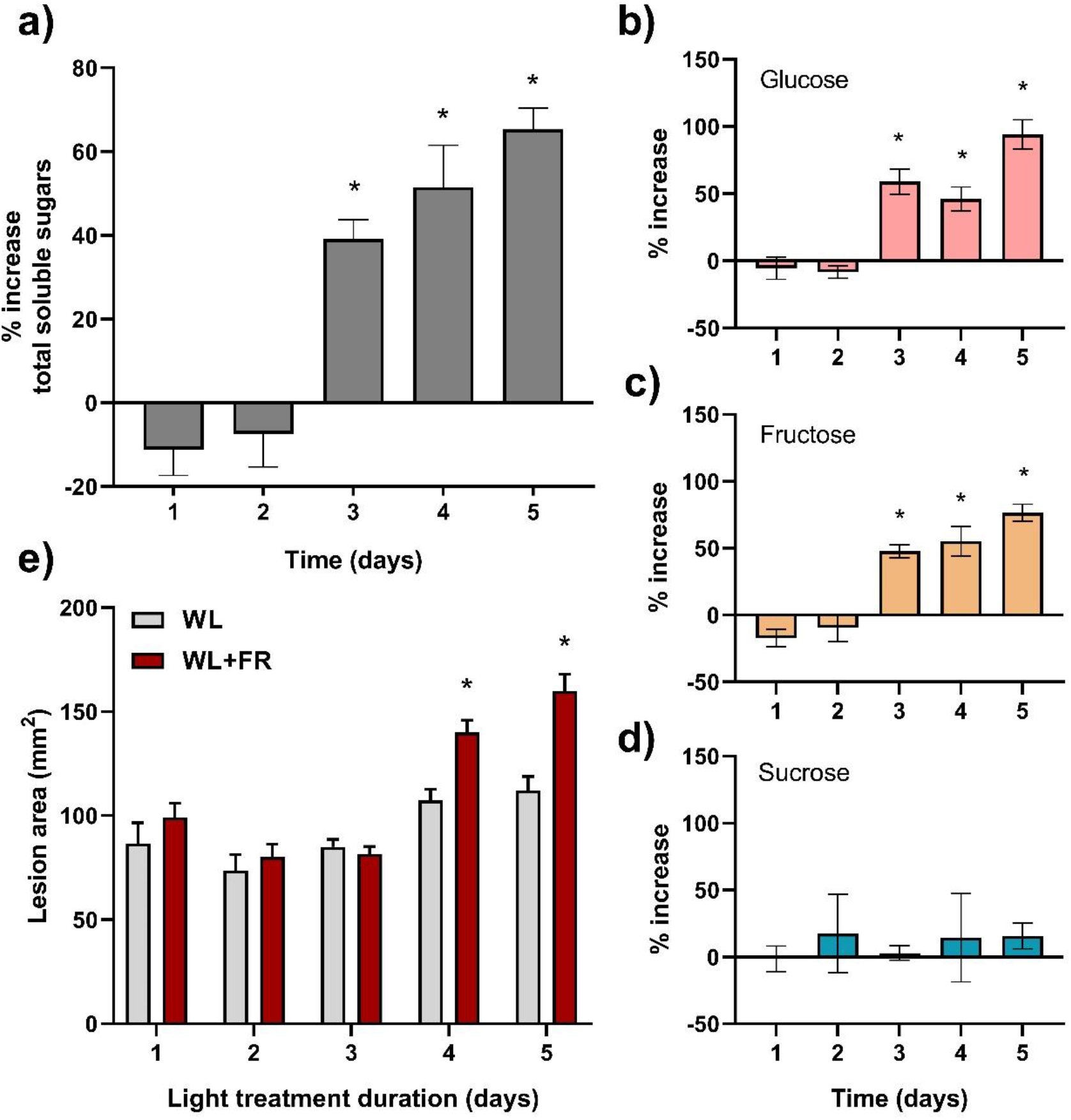
Far-red light enrichment leads to a gradual increase in soluble sugars and susceptibility to *Botrytis cinerea*. Percentage increase in **(a)** total soluble sugars, **(b)** glucose, **(c)** fructose and **(d)** sucrose levels measured in WL+FR-treated Moneymaker plants compared to WL for each timepoint. Light treatment started at 10 a.m. on day 1 (ZT = 3) and soluble sugar measurements and bioassays were performed a 3 p.m. (ZT = 8). Data correspond to mean ± SEM, n = 4 - 6 plants per treatment. The percentage increase in total soluble sugar levels was. **(e)** Disease caused by *B. cinerea* inoculated at different days after the start of WL or WL+FR treatment of three-week-old tomato leaflets. Lesion area was measured at 3 dpi. Data correspond to mean ± SEM, n = 8 plants per treatment. Asterisks represent significant differences between WL and WL+FR treatment according to Student’s t-test (p < 0.05).

### Supplemental FR promotes symptom development induced by pathogens with distinct lifestyles

The increase in soluble sugar levels observed upon WL+FR treatment in tomato plants led us to speculate that not just *B. cinerea*, but also other pathogens with different lifestyles might develop better on such leaf material since they all principally feed on sugar resources from their host. We performed bioassays on WL- and WL+FR-treated plants using *B. cinerea* (*B.c.* 05.10) as well as a pepper isolate of *B. cinerea* to test the specificity of the FR-induced susceptibility of tomato leaves in response to different *B. cinerea* isolates (Fig. **3a** and **3b**). We also tested the bacterial hemibiotroph *Pseudomonas syringae* pv. tomato DC3000 (Fig. **3c**) already described to induce more severe symptoms on Arabidopsis plants exposed to FR-enriched light conditions (de Wit *et al.*, 2013). Lastly, we tested the oomycete *Phytophthora infestans* (88069) to study the effect of supplemental FR on tomato susceptibility towards a biotrophic pathogen (Fig. **3d**). Interestingly, WL+FR-exposed tomato plants displayed significantly more severe symptoms than the WL controls to all pathogens tested (Fig. **3a-d**). Also, all pathogens showed increased growth rate when exposed to increasing concentrations of glucose *in vitro* (Fig. **3e-h**). These data indicate that the FR-induced susceptibility in tomato is not specific to *B. cinerea* as supplemental FR increased symptom development caused by an array of pathogens. We hypothesize that this phenomenon might partly be caused by the increase in soluble sugar levels upon phytochrome inactivation by WL+FR perception in tomato leaves. As an independent indication for this hypothesis, we used the *B. cinerea*-tomato pathosystem to study the effect of leaf chemical content in WL+FR-treated leaves, compared to WL-treated leaves, on *B. cinerea* mycelial growth *in vitro*. To this end, we mixed ground WL or WL+FR-treated leaf material to a minimal water-agar medium and recorded *B. cinerea* mycelial growth after three days (Fig. **S2**). As expected, the mycelium did not develop well in the water-agar control, but the addition of ground leaf material boosted the mycelial development of the fungus *in vitro* and to a higher extent on WL+FR-treated plant material than WL, even though the quantitative effect was rather small at this homogenate concentration (1 gFW ml^−1^). Additional FR was already described to increase Arabidopsis susceptibility towards *B. cinerea* and *P. syringae* via a dampening of the jasmonic acid (JA) and salicylic acid (SA) defense pathways, respectively (Cerrudo *et al.*, 2012; De Wit *et al.*, 2013). In addition, our results suggest that elevated soluble sugar levels found in tomato leaves in response to WL+FR might also promote lesion development via an accelerated growth of these pathogens in tomato leaves.

**Figure 3:**
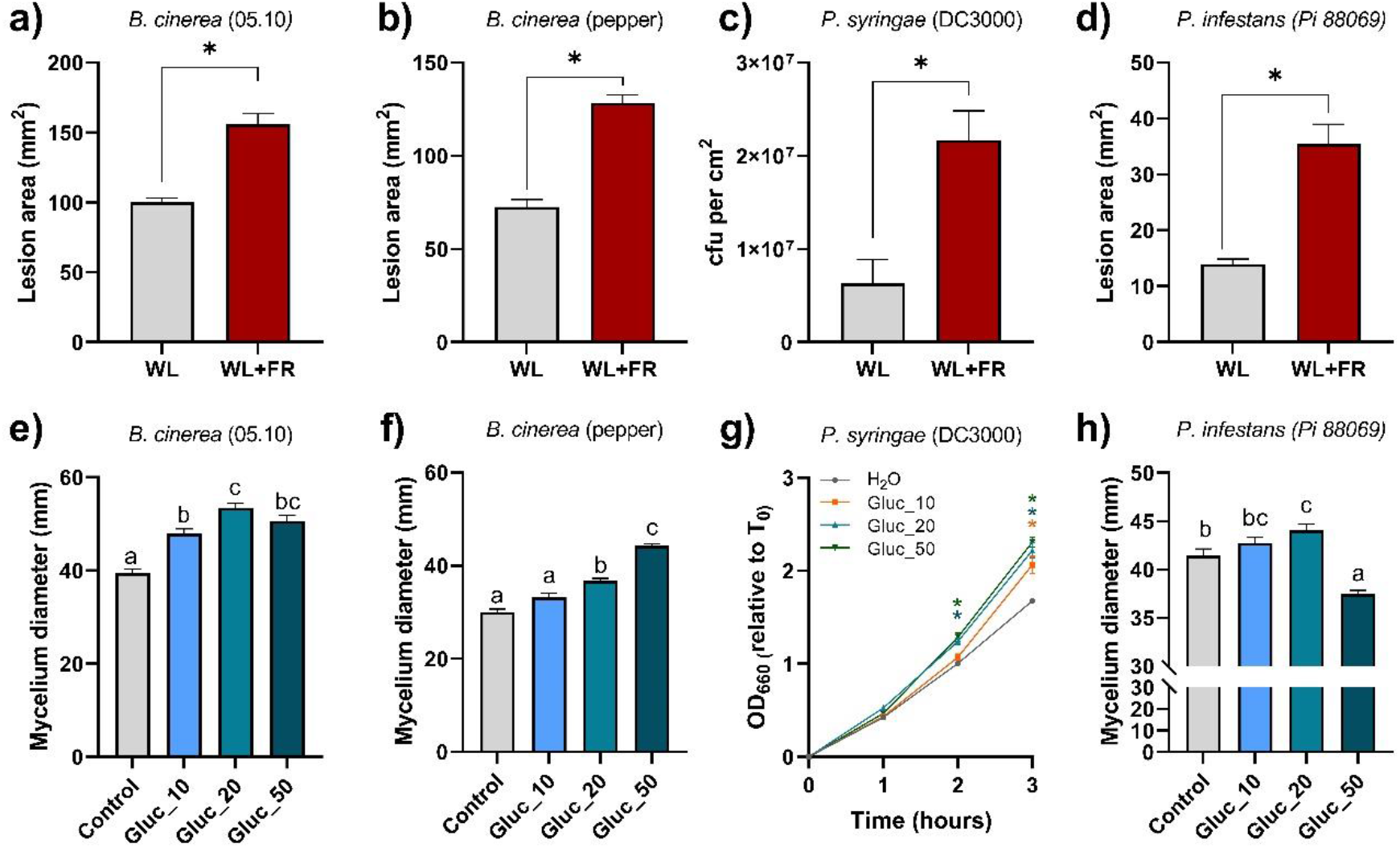
Far-red light enrichment leads to increased disease severity possibly in a sugar-dependent manner. **(a-d)** Disease rating of pathogens on three-week-old tomato plants (Moneymaker) pretreated in WL or WL+FR for five days. Pretreated leaflets were inoculated with **(a)***Botrytis cinerea* (Bc 05.10 and **(b)** another strain isolated from pepper), **(c)** the bacterial pathogen *Pseudomonas syringae* pv. tomato DC3000 and **(d)** the oomycete *Phytophthora infestans* strain 88069. **(e-h)** Pathogen growth measurements on media supplemented with increasing concentrations (0, 10, 20 or 50 g l^−1^) of glucose (Control, Gluc_10, Gluc_20 and Gluc_50). **(e and f)** Fungal growth measurement on half strength PDA-based media for *B. cinerea* strains 3 days after depositing a 5 μl droplet of a 1.5 10^6^ spores ml^−1^ solution in the center of the plates. **(g)** Bacterial growth measurements in liquid KB medium for *P*. *syringae* by measuring the OD660 every hour for three hours compared to timepoint 0. **(h)** Mycelial growth measurements for *P. infestans* on half strength V8-based media 7 days after placing a mycelium plug in the center of the plate (ø0,6 cm). n = 3 – 10. Data represent mean ± SEM. Asterisks represent significant differences according to Student’s t-test and and different letters according to ANOVA post-hoc test (p < 0.05), respectively.

### Modulation of leaf soluble sugar levels affects B. cinerea disease severity in tomato

Next, we checked whether a modulation of internal sugar levels in tomato leaves could affect *B. cinerea* lesion development. To this end, we use a detached leaflet assay allowing for passive solution uptake by the petiole prior to inoculation with the fungus (Fig. **4a**). After four days in either WL or WL+FR, tomato leaflets were detached and supplemented with either a 0.5 M glucose, 100 μM of the photosynthesis inhibitor DCMU or water as a control for 24 h prior to sugar quantification (Fig. **4b** and **4d**) or bioassays with *B. cinerea* (Fig. **4c** and **4e**). Although WL+FR-treated plants already displayed an increase in soluble sugars compared to WL-treated plants in mock conditions, the glucose solution uptake by the leaflets increased the glucose levels *in planta* after 24 h as well as the lesion area induced by *B. cinerea* (Fig. **4b** and **4c**). Consistent with this, a DCMU treatment gave opposite effects and reduced soluble sugar levels in the leaflets, which was accompanied by an increased resistance to *B. cinerea* compared to control conditions (Fig. **4d** and **4e**). These results suggest that internal sugar levels are one of the components determining disease severity in tomato leaves upon infection with *B. cinerea*. Interestingly, the lesion areas were similar between mock-treated WL+FR-exposed plants and glucose-treated WL plants even though leaf sugar levels were significantly different between both treatments (Fig. **4b** and **4c**). As the bioassays shown in figure 4 were all performed in WL conditions, the possibility of WL+FR affecting directly the pathogen growth can be ruled out. Nevertheless, we cannot exclude the effect of light quality on direct plant defense via e.g. hormone-mediated defense pathways or on the production of defense metabolites which could affect plant immunity upon WL+FR exposure (Cargnel *et al.*, 2014).

**Figure 4:**
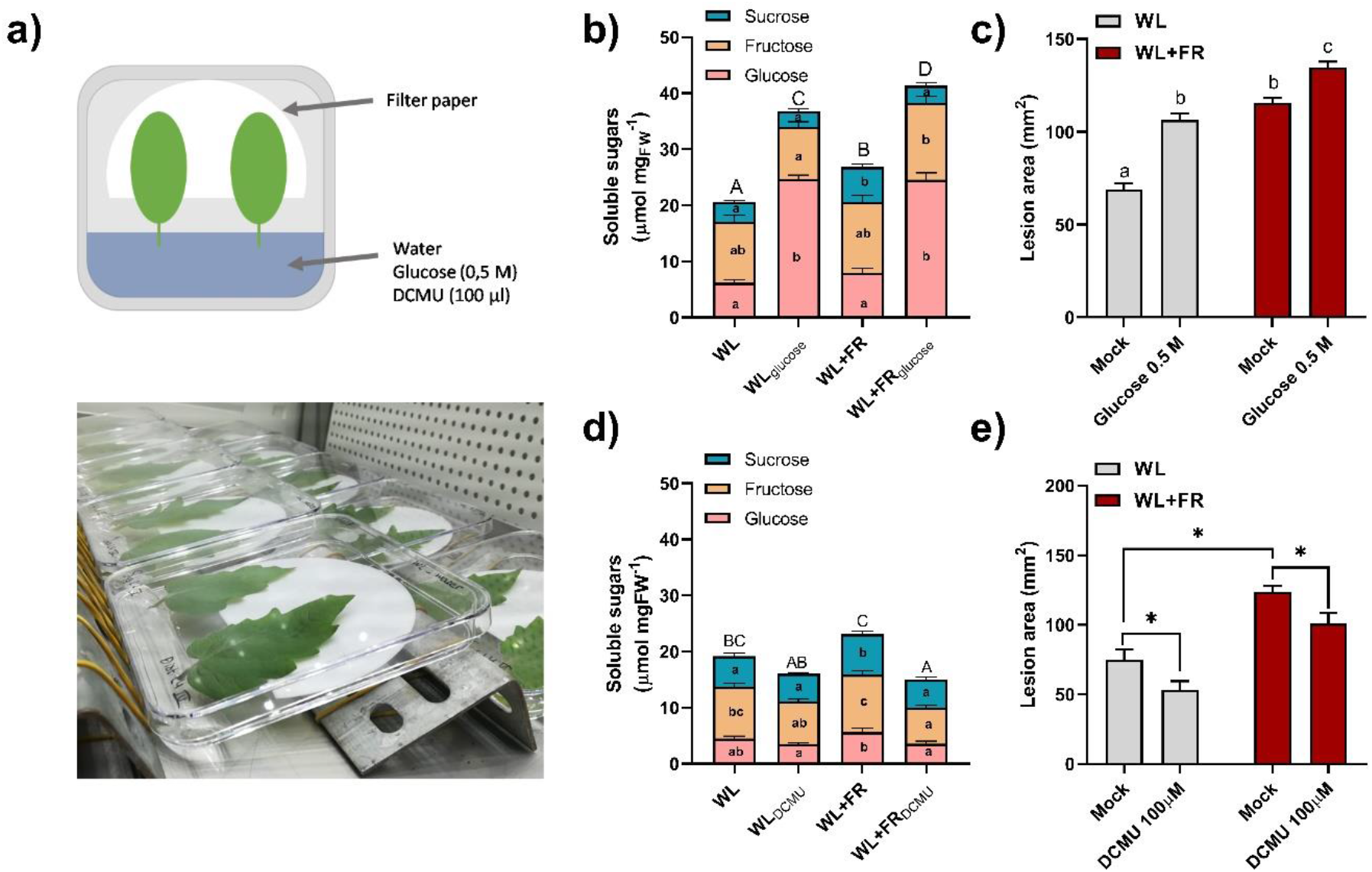
Internal soluble sugar levels in tomato leaflets dictate the severity of the symptoms induced by *Botrytis cinerea*. **(a)** Setup used to modulate soluble sugar levels in tomato detached leaflets. (**b** and **d**) Quantifications of total soluble sugars composed of glucose (pink), fructose (yellow) and sucrose (blue) fractions after five days in WL or WL+FR conditions, including a 24 h treatment with either 0.5 M glucose, 100 μM DCMU or a mock solution on detached leaflets. **(c** and **e)** Disease rating on tomato leaflets treated with glucose **(c)** or DCMU **(e)** and drop-inoculated by *B. cinerea* spores. Lesion areas were measured at 3 days post inoculation. Data show mean ± SEM, asterisks show significance difference according to Student’s t-test, letters represent significant differences according to ANOVA with Tukey’s post-hoc test (p < 0.05) with capital letters representing significance for the total soluble sugar levels (glucose + fructose + sucrose), n = 7 - 8 plants per treatment.

### Local FR enrichment influences distal disease severity to B. cinerea and soluble sugar levels

In a natural situation, FR light is rarely perceived homogeneously across the whole plant. In order to investigate whether a targeted FR enrichment (Local FR, hereafter) could have a direct influence on sugar levels and disease severity in a local and remote fashion, we designed FR lighting setups allowing for spatially restricted supplemental FR illumination (Fig. **S1**). We illuminated the top leaflet of the third oldest leaf (L3) for five days (Fig. **5a**) and observed a strong petiole elongation of the illuminated leaf compared to plants that did not receive any supplemental FR application, taken as controls (Fig. **S3**). This shows that even though FR enrichment is provided at the leaf tip (on L3), petiole elongation is still occurring throughout the leaf. Upon local FR illumination on L3, we quantified the soluble sugar content in the illuminated leaflet (L3 Loc) and in a younger distal leaflet located above, on the fourth leaf (L4 Dist) (Fig. **5a** and **5b**). After supplementing FR on L3, we observed a large increase in soluble sugars in L3 Loc but not in L4 Dist compared to the WL control for the same locations (Fig. **5b**). The increased soluble sugar levels in L3 and not in L4 were correlated with increased lesion area caused by *B. cinerea* infection in L3, but not in L4 compared to WL control plants (Fig. **5c**). Next, we swapped the treatment of the leaves: we locally exposed the top leaflet of L4 to supplemental FR (Fig. **5d**) and quantified soluble sugars and disease symptoms in this local L4 (L4 Loc), as well as in the now older, distal L3 (L3 Dist). Upon local FR illumination on L4, we observed that both L3 Dist, and L4 Loc itself exhibited higher soluble sugar content as well as increased disease symptoms compared to WL (Fig. **5e** and **5f**, respectively). Even though there might be some age-mediated differences between leaflets taken from L3 or L4, our results indicate a directional signal triggered by local FR application in the illuminated leaflet, enhancing sugar levels and susceptibility in the local leaf and in an older leaf located below, but not in younger leaves located above. The signal thus seems to move from L4 to L3 but not in the opposite direction. As the potential signal originating from the illuminated leaves is more likely to travel from young leaves towards older leaves (mainly downwards), we speculate that sugar distribution through the phloem from the younger FR-illuminated leaf to the older non-FR-illuminated leaf could have occurred. However, we cannot exclude the presence of a mobile signal, other than sugars themselves, that would be transported from the illuminated leaf towards an older leaf and trigger sugar accumulation and enhanced pathogen growth in that location. Future studies, using for example radiolabeled CO_2_ applied to the different leaf tips may help distinguish between these hypotheses.

**Figure 5:**
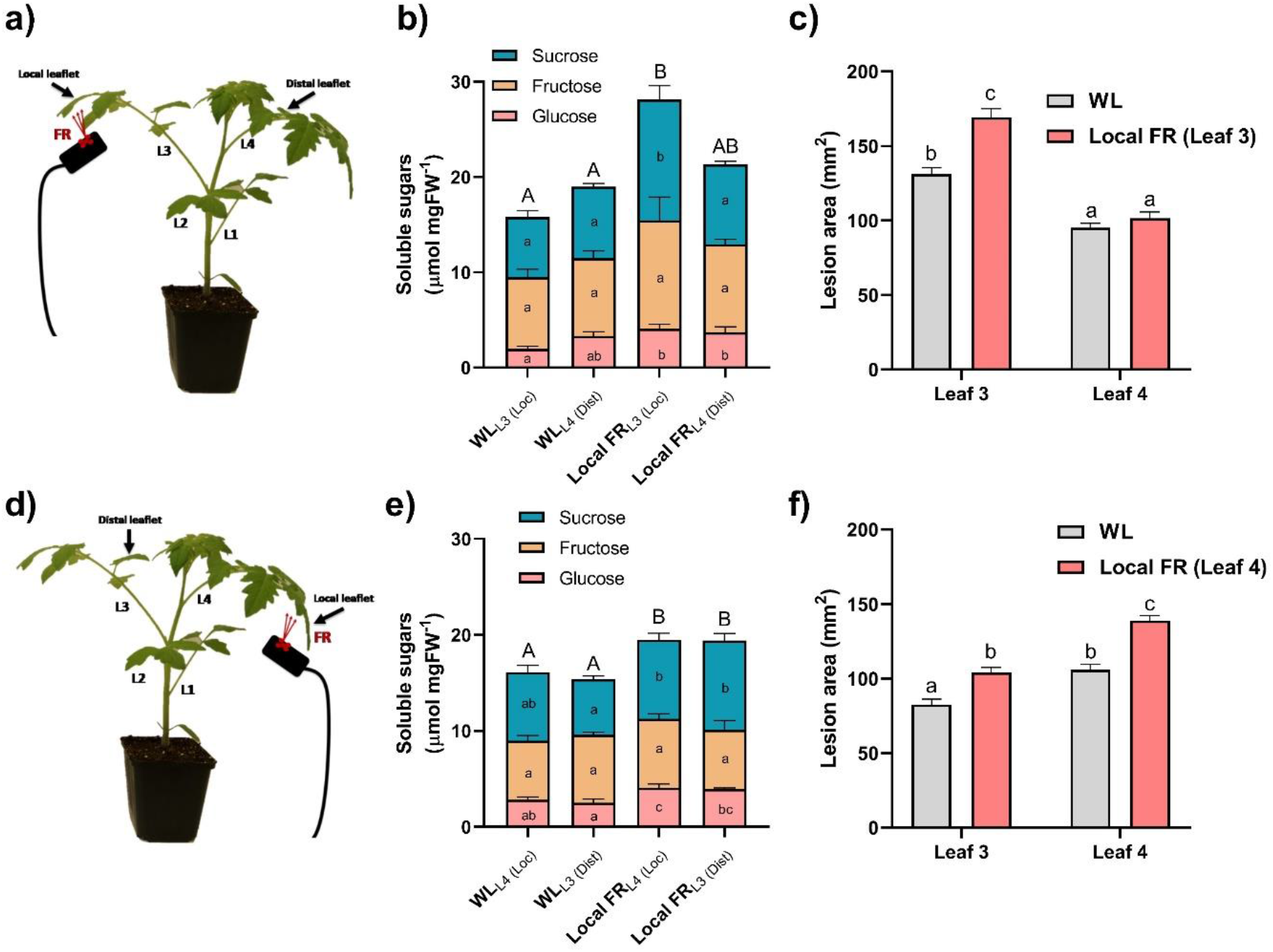
Local far-red illumination on Leaf 4 increases soluble sugar content and lesion development in both leaf 4 and leaf 3. Schematic overview of the local FR illumination procedure on leaf 3 (L3 in **a**) and on leaf 4 (L4 in **d**). Experiments were carried out on the illuminated leaflet (local: Loc) or a leaflet located on another leaf (distal: Dist). Three-week-old tomato plants were illuminated for five days with local FR (FR) or exposed to white light (WL) as a control. Soluble sugar quantifications including the glucose, fructose and sucrose fractions (**b** and **e**) as well as lesion area measurement on L3 and L4 after inoculation with *Botrytis cinerea* spores were performed at midday on day 5 (**c** and **f**). Data show mean ± SEM and different letters represent significant differences (ANOVA with Tukey’s post-hoc test, p < 0.05), n = 7 - 8 plants per treatment.

### The activation of JA signaling inhibits soluble sugar accumulation in tomato leaflets

Upon low R:FR perception in Arabidopsis or genetic inactivation of phyB in tomato, JA responsiveness is reduced (Cerrudo *et al.*, 2017; Cortés *et al.*, 2016). In addition, JA levels in plant tissue are negatively correlated to the soluble sugar levels in turn modulating disease progression in *Nicotiana attenuata* plants (Machado *et al.*, 2015). JA biosynthesis or perception mutants in tobacco displayed an increase in glucose and fructose levels but no changes in sucrose levels in leaf tissue compared to the wild-type (Machado *et al.*, 2015) similar to what we observed for the *phyB1phyB2* double mutant and WL+FR-treated tomato plants (Fig. **1b** and Fig. **2a**, respectively). To follow this up, we investigated whether the elevated soluble sugar levels observed in WL+FR-treated tomato leaves could be mediated via a JA-dependent mechanism. We exposed three-week-old tomato plants either to WL or WL+FR and sprayed the plants daily with either a 50 μM methyl jasmonate (MeJA) or a mock solution for five days prior to quantifying the soluble sugar content in the third oldest leaf (Fig. **6a**). Under WL+FR conditions the leaflets contained a higher total sugar content compared to WL conditions in the mock treatment. However, exogenous MeJA treatment dramatically reduced the levels of all soluble sugars measured (Fig. **6a**). The soluble sugar depletion induced by exogenous MeJA treatment was also associated with increased foliar resistance in WL-treated plants but not in WL+FR-treated plants possibly because of a reduced JA responsiveness in the latter (Cortés *et al.*, 2016; Fig. **6d**). In accordance, the JA biosynthesis mutant *defenseless1* (*def1*) displayed constitutively elevated glucose levels as well as increased symptom severity upon *B. cinerea* infection, compared to its wild-type background Castlemart (Fig. **6b** and **6e**). These observations confirm a connection between JA and soluble sugar levels where JA-deficient plants accumulate more soluble sugars in leaf tissue. Moreover, again we notice a correlation between elevated soluble sugar levels and enhanced pathogen infection. These data suggest that JA may play a role in soluble sugar homeostasis in tomato leaves upon WL+FR perception. Next, we attempted to rescue the *def1* mutation by providing these plants with exogenous MeJA (50 μM) every day for five days prior to quantifying soluble sugar levels or inoculating with *B. cinerea* (Fig. **6c** and **6f**). As expected, we observed a clear decrease in soluble sugar levels in *def1* mutant leaves when treated with exogenous MeJA compared to the mock (Fig. **6d**). Also, the exogenous addition of MeJA on *def1* mutants could partially rescue plant resistance to *B. cinerea* (Fig. **6e**) showing the inhibitory effect of JA signaling on soluble sugars in regulating symptom development in tomato leaves. Interestingly, the susceptibility of *def1* plants was significantly increased by the perception of supplemental FR even though the sugar levels remained unchanged (Fig. **6b** and **6e**). This observation indicates that more factors than just changes in soluble sugars levels are affected by WL+FR. This, in turn could modulate either direct plant immune responses or pathogen growth rate within plant tissue leading to more severe disease symptoms on tomato leaves. Based on these results, we hypothesize that the decreased JA responsiveness induced by WL+FR causes a soluble sugar accumulation which in turn promotes *B. cinerea* development *in planta* compared to WL conditions. Consistently, additional FR light had no effect on soluble sugar levels in the JA-deficient *def1* mutant (Fig. **6b** and **6c**). The latter experiment, however, should be interpreted cautiously since the wildtype background in which the *def1* mutant occurs (cv Castlemart) had only slight changes in soluble sugar concentrations between WL- and WL+FR-treated plants (Fig. **6b**; p = 0.056). The negative correlation between JA and soluble sugar levels was already described by Machado *et al.*, 2015. In addition, the JA biosynthesis mutant displayed an increased feeding performance by the chewing insect *Manduca sexta*. However, these caterpillars were shown to perform worse on high-sugar diets. This shows the opposite of what we describe here for WL+FR conditions, where elevated sugar levels occur together with elevated symptom development in tomato leaves by an array of pathogens (Fig. **3**). However, as the pathosystem and experimental conditions used by Machado *et al.* (2015) are hugely different from our study system it is not possible to predict whether the sugar-resistance relations in these studies contrast each other. Nevertheless, both studies describe a negative relation between JA and soluble sugar levels. Although we found a negative effect of JA on soluble sugar levels, the disease phenotypes observed do not always perfectly fit the soluble sugar levels (Fig. **6**). This observation can be explained by the effect of WL+FR on other aspect of plant immunity such as defense hormone signaling or secondary metabolite biosynthesis. As mentioned previously, glucose and JA signaling act synergistically to promote glucosinolate (GS) biosynthesis in response to *B. cinerea* in Arabidopsis (Buxdorf *et al.*, 2013; Guo *et al.*, 2013; Kliebenstein *et al.*, 2005). Even though GS are specific to Brassicaceae, it would be interesting to determine which secondary metabolites are associated with tomato resistance to *B. cinerea* and how they would be influenced by either WL+FR perception or glucose supplementation in future studies.

**Figure 6:**
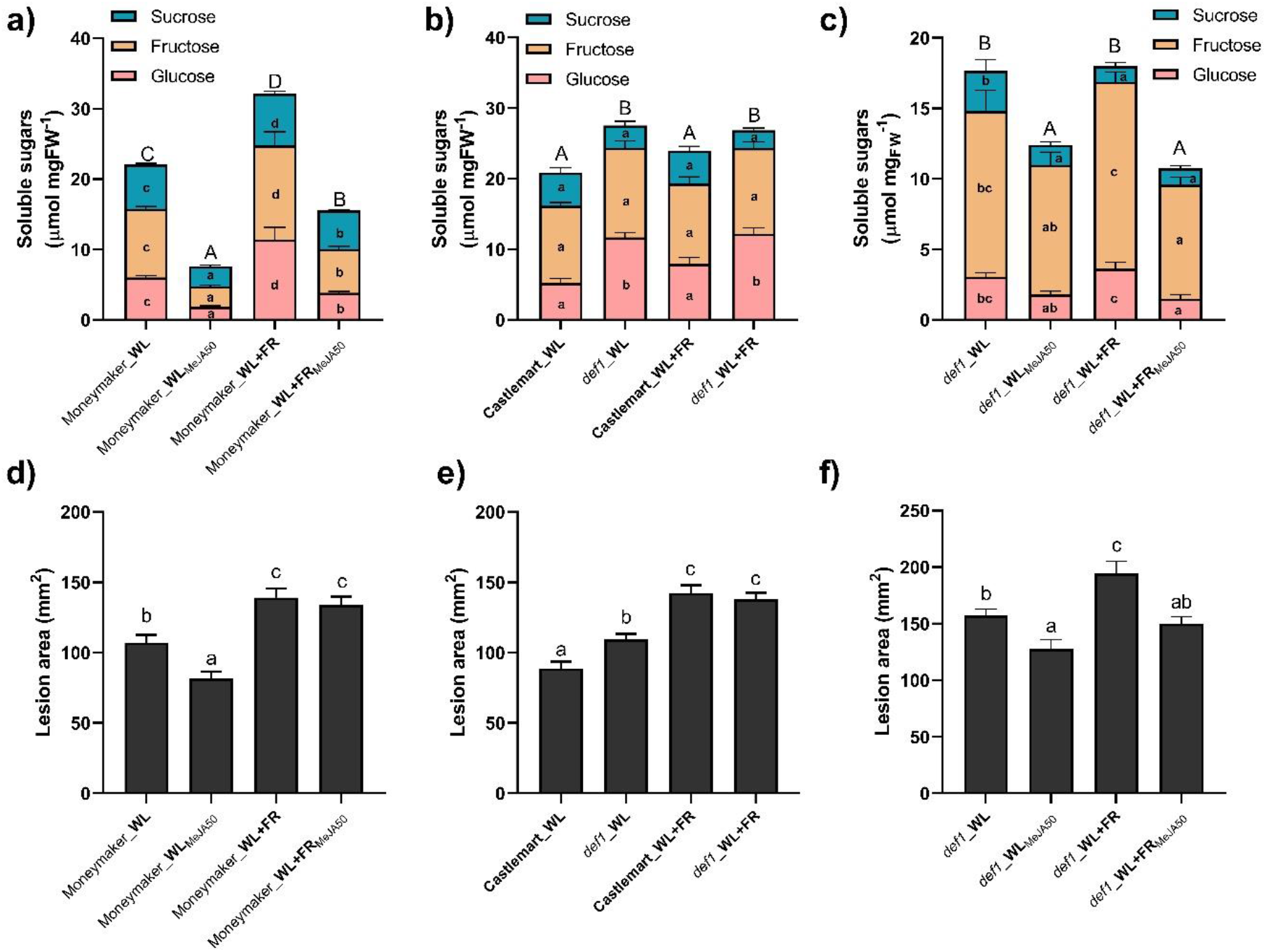
Jasmonic acid signaling inhibits sugar accumulation in tomato associated with increased lesion development. **(a-c)** Leaf total soluble sugar levels including glucose (pink), fructose (yellow) and sucrose (blue) fractions as well as **(d-f)** lesion area induced by *B. cinerea* on detached leaflets at 3 dpi in **(a and d)** Moneymaker plants exposed for five days to WL or WL+FR conditions and treated daily with 50 μM MeJA or a mock solution, **(b and e)** in tomato cv. Castlemart plants and the JA biosynthesis mutants *def1* (*defenseless 1*) exposed to WL or WL+FR conditions for five days or **(c and f)** in the *def1* mutant treated for five days in WL and WL+FR and treated daily with a 50 μM MeJA or a mock solution. Different letters represent significant differences according to ANOVA, Tukey’s post-hoc test (p < 0.05). Data represent mean ± SEM, n = 3 - 8 plants per treatment.

## Conclusion and perspectives

Our results show that a decrease in JA responsiveness previously observed in tomato (Cortes *et al.*, 2016) coupled with elevated soluble sugar levels promotes pathogen growth in plant tissue leading to increased lesion development. We demonstrate that supplemental FR, either applied to the whole plant or on a targeted leaflet, interferes with soluble sugar homeostasis in tomato leaves in a local and remote fashion thereby affecting pathogen growth capacity in plant tissue. We also show evidence about the role of JA in the regulation of soluble sugar levels upon WL+FR exposure. Although the soluble sugar status of the plants alone does not always predict the susceptibility phenotype upon infection by *B. cinerea*, we show that the modulation of soluble sugar levels in tomato leaves is mostly based on a JA-dependent mechanism and is overall associated with severity of disease symptoms.

In the context of global climate change and the simultaneously increasing global food demand, novel agricultural approaches are needed to increase crop yield under challenging environmental conditions. One option to do so is to grow plants closer together, whilst maintaining yield and resilience. However, we show that neighbor detection via a reduction in the R:FR ratio elevates foliar susceptibility to pathogenic microorganisms via a rapid increase in soluble sugar levels. Nevertheless, in greenhouse systems, applying supplemental light using LEDs is relatively straightforward. Assuming that supplemental red LEDs would be applied as inter-lighting to counteract the naturally occurring drop in R:FR due to the absorption of red light by the crop stand it might be possible to maintain plant resistance even at high density. In a dense plant canopy, the elevation of soluble sugars in WL+FR could possibly be controlled by increasing the R:FR ratio in the middle of the canopy in order to reduce the sugar-mediated susceptibility that occurs systemically in older, but not younger leaves than the FR-exposed ones (Fig. **5**). Higher in the canopy, there is still ample direct light from top lamps and outside to keep R:FR relatively high. Overall, increasing plant resistance at high density by using LED lighting seems possible and could partly solve the FR-induced susceptibility in tomato towards pathogens. Also, as sugar supplies are beneficial to most pathogens, we believe that balancing soluble sugar levels *in planta* by smart LED lighting plans would rescue tomato susceptibility to other pathogens than *B. cinerea* as well. Future studies on timing and spatial distribution of light quality all over a mature tomato stand are needed to understand if these positive effects of FR-enriched light can be achieved without having the negative effects on tomato pathogen resistance.

## Acknowledgements

We thank Hans van Veen and Putri Prasetyaningrum for help with soluble sugar quantifications and the LED it Be 50% consortium, Jesse Küpers and Sjon Hartman for helpful discussions. This work was funded by the Dutch Research Council, TTW Perspectief grant nr 14125 (LED it Be 50%) with contributions from Signify, LTO Glaskracht and WUR Greenhouse Horticulture.

## Author contributions

S.C. and R.P. designed the study and experiments with additional input from K.K. S.G., E.S., P-O.B. and S.C. carried out the experiments. S.C. drafted the article and designed the figures. R.P. and S.C. discussed results, R.P., S.C. and S.W. wrote the final manuscript, K.K. commented on the final draft.

## Supplemental data

**Figure S1:**
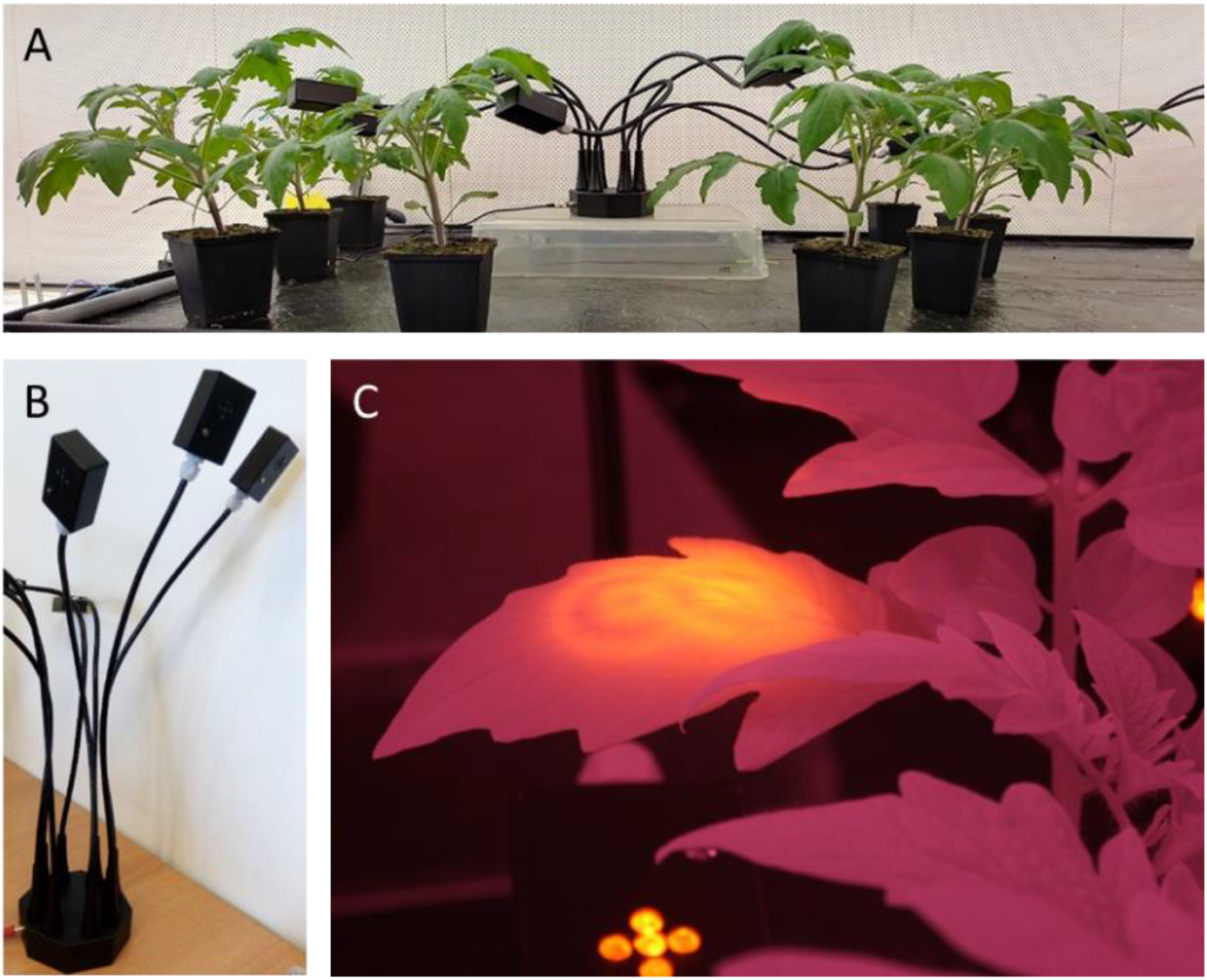
Local FR illumination setup. The top leaflet of leaf n°3 was illuminated from the bottom (**A**) by using FR LEDs attached to flexible arms (**B**). Infrared pictures showing FR illumination of the center of the lamina (**C**).

**Figure S2:**
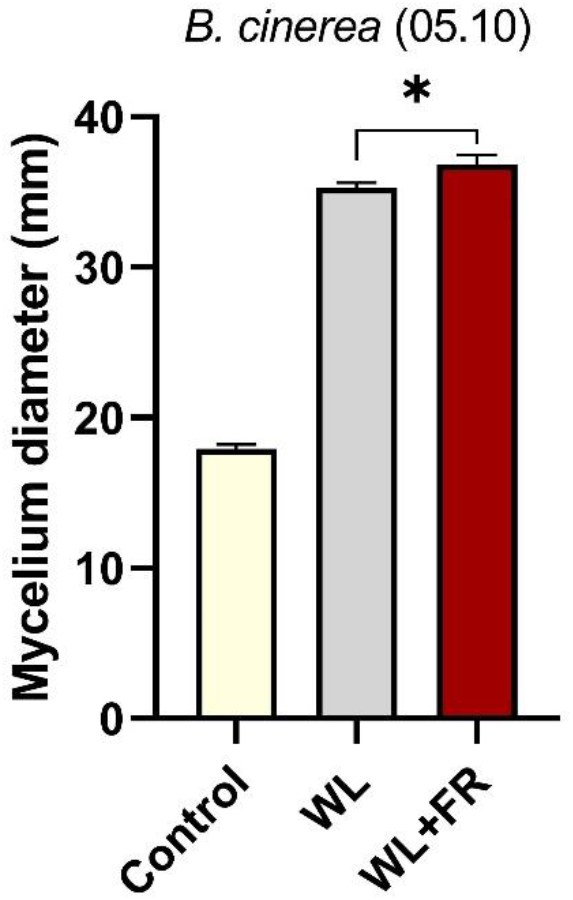
*Botrytis cinerea* mycelial growth is enhanced by elevated soluble sugar concentrations *in planta.* Mycelium diameter growth measurement after three days on water-based agar media supplemented with either water (Control) WL-treated or WL+FR-treated leaf homogenate (WL and WL+FR, respectively). Data represent mean ± SEM and asterisk represent significant difference (Student’s t-test, p < 0.05, n = 10).

**Figure S3:**
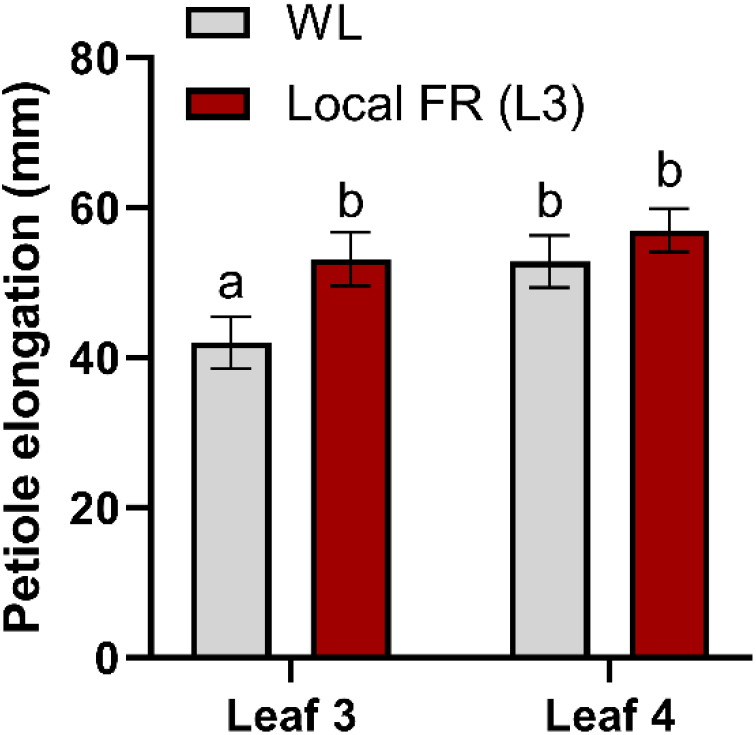
Local FR enrichment triggers petiole elongation. Petiole elongation measurements of the third and fourth leaf of three-week-old tomato plants after five days of local FR illumination on the top leaflet of the third leaf. Measurements were performed at the start and after five days of WL or local FR treatment light treatment (grey and red bars, respectively). Data represent three replicates merged together (n = 6 - 8 plants). Plotted data correspond to mean ± SEM. Different letters represent significant differences (ANOVA Tukey’s post-hoc test, p < 0,05).

## References

Ballaré, C.L. 2014. Light regulation of plant defense. Annual Review of Plant Biology 65: 335–363.

Ballaré, C.L., Pierik, R. 2017. The shade-avoidance syndrome: Multiple signals and ecological consequences. Plant Cell and Environement 40: 2530–2543.

Buxdorf, K., Yaffe, H., Barda, O., Levy, M. 2013. The Effects of Glucosinolates and Their Breakdown Products on Necrotrophic Fungi. PLoS One 8: e70771.

Cargnel, M.D., Demkura, P. V., Ballaré, C.L. 2014. Linking phytochrome to plant immunity: low red: Far-red ratios increase Arabidopsis susceptibility to *Botrytis cinerea* by reducing the biosynthesis of indolic glucosinolates and camalexin. New Phytologist 204: 342–354.

Caten, C.E., Jinks, J.L. 1968. Spontaneous variability of single isolates of *Phytophthora infestans*. I. Cultural variation. Canadian Journal of Botany 46: 329–348.

Cerrudo, I., Caliri-Ortiz, M.E., Keller, M.M., Degano, M.E., Demkura, P. V., Ballaré, C.L. 2017. Exploring growth-defence trade-offs in Arabidopsis: phytochrome B inactivation requires JAZ10 to suppress plant immunity but not to trigger shade-avoidance responses. Plant Cell and Environement 40: 635–644.

Cerrudo, I., Keller, M.M., Cargnel, M.D., Demkura, P. V, de Wit, M., Patitucci, M.S., Pierik, R., Pieterse, C.M.J., Ballaré, C.L. 2012. Low red/far-red ratios reduce Arabidopsis resistance to Botrytis cinerea and jasmonate responses via a COI1-JAZ10-dependent, salicylic acid-independent mechanism. Plant Physiology 158: 2042–52.

Chen, L.Q., Hou, B.H., Lalonde, S., Takanaga, H., Hartung, M.L., Qu, X.Q., Guo, W.J., Kim, J.G., Underwood, W., Chaudhuri, B., Chermak, D., Antony, G., White, F.F., Somerville, S.C., Mudgett, M.B., Frommer, W.B. 2010. Sugar transporters for intercellular exchange and nutrition of pathogens. Nature 468: 527–532.

Chong, J., Piron, M.C., Meyer, S., Merdinoglu, D., Bertsch, C., Mestre, P. 2014. The SWEET family of sugar transporters in grapevine: VvSWEET4 is involved in the interaction with *Botrytis cinerea*. Journal of Experimental Botany 65: 6589–6601.

Cortés, L.E., Weldegergis, B.T., Boccalandro, H.E., Dicke, M., Ballaré, C.L. 2016. Trading direct for indirect defense? Phytochrome B inactivation in tomato attenuates direct anti-herbivore defenses whilst enhancing volatile-mediated attraction of predators. New Phytologist 212: 1057–1071.

De Wit, M., George, G.M., Ince, Y.Ç., Dankwa-Egli, B., Hersch, M., Zeeman, S.C., Fankhauser, C. 2018. Changes in resource partitioning between and within organs support growth adjustment to neighbor proximity in Brassicaceae seedlings. Proceedings of the National Academy of Sciences of the United States of America 115: E9953–E9961.

De Wit, M., Spoel, S.H., Sanchez-Perez, G.F., Gommers, C.M.M., Pieterse, C.M.J., Voesenek, L.A.C.J., Pierik, R. 2013. Perception of low red: Far-red ratio compromises both salicylic acid- and jasmonic acid-dependent pathogen defences in Arabidopsis. The Plant Journal 75: 90–103.

Denby, K.J., Kumar, P., Kliebenstein, D.J. 2004. Identification of *Botrytis cinerea* susceptibility loci in *Arabidopsis thaliana*. The Plant Journal 38: 473–486.

Doidy, J., Grace, E., Kühn, C., Simon-Plas, F., Casieri, L., Wipf, D. 2012. Sugar transporters in plants and in their interactions with fungi. Trends in Plant Science 17: 413–422.

Fernández-Milmanda, G.L., Crocco, C.D., Reichelt, M., Mazza, C.A., Köllner, T.G., Zhang, T., Cargnel, M.D., Lichy, M.Z., Fiorucci, A.S., Fankhauser, C., Koo, A.J., Austin, A.T., Gershenzon, J., Ballaré, C.L. 2020. A light-dependent molecular link between competition cues and defence responses in plants. Nature Plants 6: 223–230.

Franklin, K.A. 2008. Shade avoidance. New Phytologist 179: 930–944.

Grant, J.J., Chini, A., Basu, D., Loake, G.J. 2003. Targeted Activation Tagging of the Arabidopsis NBS-LRR gene, ADR1, Conveys Resistance to Virulent Pathogens. Molecular Plant-Microbe Interactions 16: 669–680.

Guo, R., Shen, W., Qian, H., Zhang, M., Liu, L., Wang, Q. 2013. Jasmonic acid and glucose synergistically modulate the accumulation of glucosinolates in *Arabidopsis thaliana*. Journal of Experimental Botany 64: 5707–5719.

Hornitschek, P., Kohnen, M. V., Lorrain, S., Rougemont, J., Ljung, K., López-Vidriero, I., Franco-Zorrilla, J.M., Solano, R., Trevisan, M., Pradervand, S., Xenarios, I., Fankhauser, C. 2012. Phytochrome interacting factors 4 and 5 control seedling growth in changing light conditions by directly controlling auxin signaling. Plant Journal 71: 699–711.

Izaguirre, M.M., Mazza, C.A., Biondini, M., Baldwin, I.T., Ballaré, C.L. 2006. Remote sensing of future competitors: Impacts on plants defenses. Proceedings of the National Academy of Sciences of the United States of America 103: 7170–7174.

Ji, Y., Ouzounis, T., Courbier, S., Kaiser, E., Nguyen, P.T., Schouten, H.J., Visser, R.G.F., Pierik, R., Marcelis, L.F.M., Heuvelink, E. 2019. Far-red radiation increases dry mass partitioning to fruits but reduces *Botrytis cinerea* resistance in tomato. Environmental and Experimental Botany 168: 103889.

Kamoun, S., Van West, P., De Jong, A.J., De Groot, K.E., Vleeshouwers, V.G.A.A., Govers, F. 1997. A gene encoding a protein elicitor of *Phytophthora infestans* is downregulated during infection of potato. Molecular Plant-Microbe Interactions 10: 13–20.

Kasperbauer, M.J., Tso, T.C., Sorokin, T.P. 1970. Effects of end-of-day red and far-red radiation on free sugars, organic acids and amino acids of tobacco. Phytochemistry 9: 2091–2095.

King, E.O., Ward, M.K., Raney, D.E. 1954. Two simple media for the demonstration of pyocyanin and fluorescin. The Journal of Laboratory and Clinical Medicine 44: 301–307.

Kliebenstein, D.J., Rowe, H.C., Denby, K.J. 2005. Secondary metabolites influence Arabidopsis/Botrytis interactions: Variation in host production and pathogen sensitivity. Plant Journal 44: 25–36.

Lapin, D., Van den Ackerveken, G. 2013. Susceptibility to plant disease: More than a failure of host immunity. Trends in Plant Science 18: 546–554.

Lecompte, F., Nicot, P.C., Ripoll, J., Abro, M.A., Raimbault, A.K., Lopez-Lauri, F., Bertin, N. 2017. Reduced susceptibility of tomato stem to the necrotrophic fungus *Botrytis cinerea* is associated with a specific adjustment of fructose content in the host sugar pool. Annals of Botany 119: 931–943.

Leone, M., Keller, M.M., Cerrudo, I., Ballaré, C.L. 2014. To grow or defend? Low red: Far-red ratios reduce jasmonate sensitivity in Arabidopsis seedlings by promoting DELLA degradation and increasing JAZ10 stability. New Phytologist 204: 355–367.

Lercari, B. 1982. The effect of far‐red light on the photoperiodic regulation of carbohydrate accumulation in Allium cepa L. Physiologia Plantarum 54: 475–479.

Li, K., Yu, R., Fan, L.M., Wei, N., Chen, H., Deng, X.W. 2016. DELLA-mediated PIF degradation contributes to coordination of light and gibberellin signalling in Arabidopsis. Nature Communications 7: 11868.

Li, L., Ljung, K., Breton, G., Schmitz, R.J., Pruneda-Paz, J., Cowing-Zitron, C., Cole, B.J., Ivans, L.J., Pedmale, U. V., Jung, H.S., Ecker, J.R., Kay, S.A., Chory, J. 2012. Linking photoreceptor excitation to changes in plant architecture. Genes and Development 26: 785–790.

Machado, R.A.R., Arce, C.C.M., Ferrieri, A.P., Baldwin, I.T., Erb, M. 2015. Jasmonate-dependent depletion of soluble sugars compromises plant resistance to *Manduca sexta*. New Phytologist 207: 91–105.

Moghaddam, M.R.B., Van Den Ende, W. 2012. Sugars and plant innate immunity. Journal of Experimental Botany 63: 3989–3998.

Pantazopoulou, C.K., Bongers, F.J., Küpers, J.J., Reinen, E., Das, D., Evers, J.B., Anten, N.P.R., Pierik, R. 2017. Neighbor detection at the leaf tip adaptively regulates upward leaf movement through spatial auxin dynamics. Proceedings of the National Academy of Sciences 114: 7450–7455.

Pieterse, C. M. J., Pierik, R., Van Wees, S.C.M. 2014. Different shades of JAZ during plant growth and defense. New Phytologist 204: 261–264.

Sager, J.C., Smith, W.O., Edwards, J.L., Cyr, K.L. 1988. Photosynthetic efficiency and phytochrome photoequilibria determination using spectral data. Transactions of the American Society of Agricultural Engineers 31: 1882–1889.

Van Kan, J.A.L., Van’t Klooster, J.W., Wagemakers, C.A.M., Dees, D.C.T., Van Der Vlugt-Bergmans, C.J.B. 1997. Cutinase A of *Botrytis cinerea* is expressed, but not essential, during penetration of gerbera and tomato. Molecular Plant-Microbe Interactions 10: 30–38.

Van Wees, S.C.M., Van Pelt, J.A., Bakker, P.A.H.M., Pieterse, C.M.J. 2013. Bioassays for assessing jasmonate-dependent defenses triggered by pathogens, herbivorous insects, or beneficial rhizobacteria. Methods in Molecular Biology 1011: 35–49.

Whalen, M.C., Innes, R.W., Bent, A.F., Staskawicz, B.J. 1991. Identification of *Pseudomonas syringae* pathogens of Arabidopsis and a bacterial locus determining avirulence on both Arabidopsis and soybean. The Plant Cell 3: 49–59.

Xiong, J., Patil, G.G., Moe, R., Torre, S. 2011. Effects of diurnal temperature alternations and light quality on growth, morphogenesis and carbohydrate content of *Cucumis sativus* L. Scientia Horticulturae 128: 54–60.

Yang, D., Seaton, D.D., Krahmer, J., Halliday, K.J. 2016. Photoreceptor effects on plant biomass, resource allocation, and metabolic state. Proceedings of the National Academy of Sciences 113: 7667–7672.

